# Exploring the genetic architecture underlying dietary fiber content in Colombian Andean blueberry (*Vaccinium meridionale* Swartz)

**DOI:** 10.64898/2026.02.19.706934

**Authors:** Ginna Patricia Velasco Anacona, Angie Carolina Guevara Correa, Carlos Eduardo Narváez Cuenca, Teresa Mosquera Vásquez, Johana Carolina Soto Sedano

## Abstract

Dietary fiber composition is a major determinant of fruit nutritional quality, yet its genetic basis remains poorly characterized in wild Vaccinium species. Here, we combined extensive phenotyping with a genome-wide association study (GWAS) to dissect the genetic control of dietary fiber traits in Colombian agraz (*Vaccinium meridionale* Swartz). Total dietary fiber (TDF), insoluble dietary fiber (IDF), and soluble dietary fiber (SDF), the SDF/IDF ratio, and maturity index (MI) were quantified in fruits from 119 genotypes, representing the most comprehensive evaluation of dietary fiber fractions in fresh Vaccinium fruit to date. GWAS mapped this phenotypic diversity to 24 QTLs distributed across 15 chromosomes, revealing a polygenic architecture underlying fiber-related trait. A TDF QTL (Chr41:26883013) directly co-localized with *VaccDscaff31-augustus-gene-268.33*, a 7-deoxyloganetin glucosyltransferase, embedded within a glycosyltransferase-rich LD block. IDF variation was associated with *VaccDscaff33-processed-gene-116.2* (pectin methylesterase 15) while the SDF/IDF ratio co-localized with *VaccDscaff55-augustus-gene-9.30*, encoding a xyloglucan endotransglucosylase/hydrolase. Together, the integration of high-resolution phenotyping with QTL mapping connects natural variation in dietary fiber content and composition to specific biosynthetic, remodeling, and regulatory pathways, providing actionable molecular targets for marker-assisted and genomic selection aimed at improving nutritional quality, texture, and processing traits in Vaccinium breeding programs.

## Introduction

Blueberries are widely recognized as a health-protective fruit, primarily due to their bioactive compounds, especially flavonoids (such as anthocyanins), flavonols (such as quercetin [1]glycosides), phenolic acids (such as chlorogenic acid) [1], and dietary fiber [2]. The Colombian blueberry, known as agraz (*Vaccinium meridionale* Swartz), is a wild species belonging to the Ericaceae family, that grows spontaneously in the mountainous regions of Colombia, Ecuador, Peru, Venezuela, and Jamaica [3]. Its fruit is an edible berry with an acidic and astringent taste, valued by consumers who demand natural sources of beneficial compounds. It is consumed fresh, dehydrated or in preparations such as wines, nectars, and jams [4]. Previous studies indicate that agraz fruits contain a variety of polyphenolic compounds and dietary fiber, which are associated with significant bioactive properties, such as high antioxidant capacity [5], antiproliferative, and cytotoxic effects against cancer cells [6], antimicrobial activity, and protection against colorectal cancer [7,8].

Dietary fiber is an important component in foods since it is related to human health. A recent meta-analysis including 17,155,277 individuals reported that high dietary fiber intake is associated with a reduced risk of several chronic diseases, particularly cardiovascular disease mortality, pancreatic cancer, and diverticular disease [9]. Although we have recently reported information on the chemical and genetic architecture of polyphenolic composition in Colombian *V. meridionale* fruits [10], information regarding the variability of dietary fiber fractions in this species remains limited. Moreover, studies evaluating dietary fiber composition across large genotype collections are scarce not only for agraz but also for other *Vaccinium* species.

Dietary fiber in fruits is a complex and dynamic trait primarily determined by the composition and structural organization of plant cell wall polysaccharides [11]. It is mainly composed of cellulose, hemicellulosic polysacharides, lignin, and pectins, which together define the mechanical properties, texture, and nutritional quality of fleshy fruits [11].

From a nutritional perspective, dietary fiber is conventionally classified into insoluble dietary fiber (IDF) and soluble dietary fiber (SPF). IDF is largely associated with cellulose, lignin, and hemicellulosic polysaccharides [12], while SDF, is dominated by pectins and hemicellulosic polysaccharides [13]. Hemicellulosic polysaccharides are present in both fractions; however, they differ markedly in structural complexity, branching patterns, and association with the plant cell wall matrix. In the IDF fraction, hemicelluloses are predominantly composed of structurally integrated polymers that are tightly bound to cellulose microfibrils and lignin [12]. These typically include less branched xylans and structural mannans or glucomannans, which contribute to cell wall rigidity and exhibit limited water solubility [11]. In contrast, the hemicellulosic polysaccharides found in the SDF fraction are generally more loosely associated with the cell wall or are readily extractable [12]. These polymers are often characterized by higher degrees of branching and structural heterogeneity, including arabinoxylans with extensive arabinose substitutions, xyloglucans, and mixed-linkage β-glucans, which enhance hydration properties and solubility [13].

Understanding the structural differences between fiber fractions requires consideration of the biosynthetic pathways that determine cell wall polymer assembly and remodeling. Cellulose is synthesized by *CesA* genes, which encode the subunits of the cellulose synthase enzyme complex, visible in the plasma membrane [14]. Hemicelluloses comprise a diverse group of heterogeneous polysaccharides that function as cross-linking components between cellulose microfibrils in the primary cell wall, including heteromannans, xyloglucans, heteroxylans, and mixed-linkage glucans [15]. Their biosynthesis is mediated predominantly by glycosyltransferases (GTs), including uridine 5′-diphospho-glucuronosyltransferases (UDP-glycosyltransferases). The synthesis of the xyloglucan backbone is catalyzed by members of the cellulose synthase-like (*CSL*) family, enzymes belonging to the GT2-CSL superfamily [15]. Within this group, distinct *CSL* clades specialize in the synthesis of different hemicellulosic polymers: CSL-C proteins are involved in xyloglucan chain elongation, CSL-A members participate in the biosynthesis of mannan and glucomannan backbones, and CSL-F enzymes contribute to mixed-linkage glucan formation [15]. In contrast, the assembly of xylan backbones is mediated by glycosyltransferases of the GT43 family, which exhibit β-1,4-xylosyltransferase activity [16]. Hemicellulose remodeling involves enzymes such as xyloglucan endotransglucosylase/hydrolases (XTHs), which modify xyloglucan chain length and connectivity [17], as well as β-xylosidases and α-L-arabinofuranosidases, which participate in the degradation and restructuring of hemicellulosic polysaccharides during cell wall remodeling [18,19].

In addition to polysaccharide components, the insoluble fiber fraction also includes lignin, a complex phenolic polymer that reinforces secondary cell walls and contributes to tissue rigidity. Lignin biosynthesis occurs through the phenylpropanoid pathway and is mediated by the coordinated action of enzymes such as cinnamoyl-CoA reductase (CCR), 4-cinnamoyl-CoA ligase (4CL), shikimate *O*-hydroxycinnamoyltransferase (HCT), and peroxidases (PODs), which contribute to secondary wall strengthening and tissue rigidity [20]. In *Vaccinium corymbosum*, genome-wide analyses identified 92 caffeic acid *O*-methyltransferase (COMT) genes, with specific members such as *VcCOMT40* and *VcCOMT92* showing a direct relationship with lignin accumulation during fruit development, underscoring their relevance for fruit firmness and shelf life [21].

Finally, pectin, a heterogeneous group of complex polymers composed mainly of homogalacturonan (HG) and rhamnogalacturonan I and II (RG-I and RG-II) [22,23]. Homogalacturonan is the most abundant pectic domain and consists of linear chains of (1→4)-α-D-galacturonic acid residues, whose backbone is synthesized in the Golgi apparatus by galacturonosyltransferases (GAUTs) [24]. Following biosynthesis, pectin undergoes extensive remodeling mediated by enzymes such as pectin methylesterases (PMEs) and polygalacturonases (PGs), which regulate the degree of methylesterification, depolymerization, and interactions with other cell wall polymers, thereby modulating pectin solubility and the mechanical properties of the cell wall [25].

Based on the above considerations, we hypothesize that there are genetic mechanisms underlying the natural variability in the dietary fiber in *V. meridionale*. Accordingly, this study aimed to quantify IDF, SDF, total dietary fiber (TDF), and the SDF/IDF × 100 ratio in a large collection of ripe agraz fruit genotypes collected in rural areas of the Colombian Departments of Santander, Boyacá, and Cundinamarca. In addition, a genome-wide association study (GWAS) was conducted in a representative agraz subpopulation to identify quantitative trait loci (QTL) potentially associated with dietary fiber traits. The results of this study provide deeper insight into the genetic mechanisms underlying dietary fiber content in agraz and establish a scientific foundation for future breeding strategies aimed at improving fiber-related nutritional quality. Moreover, these findings highlight the importance of advancing research in wild and underutilized crops with high nutritional and commercial potential, emphasizing their value as genetic reservoirs for crop improvement and sustainable agriculture.

## Materials and methods

### Fruit samples

A previously reported collection of 118 fruit genotypes sampled from rural areas of Santander, Boyacá, and Cundinamarca (Colombia) was established as a genetically diverse *V. meridionale* population [10,26]. Also, a control commercial *V. meridionale* sample was analyzed [10] to complete a collection of 119 fruit material (S1 Table).

### Dietary fiber content

IDF and SDF contents were measured by the enzymatic-gravimetric method as described in the AOAC 985.29 and AOAC 991.43 methods [27] in the freeze-dried-ground samples. Both components (IDF and SDF) were experimentally found by performing three replications. Corrections because of ash and protein contents were performed for both, SDF and ISF contents. Additionally, TDF content was calculated as the sum of SDF and IDF. For sample digestion, 0.5 g of each freeze-dried sample was weighed and subjected to treatments with α-amylase, protease, and amyloglucosidase, adjusting the pH and temperature at each stage to facilitate the digestion of specific components. Dietary fiber kit was purchased from Sigma-Aldrich (Darmstad, Germany). IDF content was determined by filtering the resulting digestate, followed by sequential washing of the retained residue with water, 78% aqueous ethanol, 95% aqueous ethanol, and acetone; the residue was then dried at 105 °C until constant weight. SDF was determined by mixing the filtrate with 95% (*v/v*) aqueous ethanol at a ratio of 1:4 (filtrate: aqueous ethanol, *v/v*). The mixture was allowed to stand overnight at 4 °C to precipitate soluble fiber components. The precipitate was recovered by filtration and sequentially washed with 78% (*v/v*) aqueous ethanol, 95% aqueous ethanol, and acetone. The recovered SDF was then dried at 105 °C to constant weight. Both types of fiber contents were corrected for protein and ash content. Method reproducibility was assessed by performing technical replicates for selected genotypes. Results are expressed as mean values ± standard deviation, reported on both a dry weight (DW) and fresh weight (FW) basis.

To verify if the measured dietary fiber variables were correlated with the maturity state of the collected fruit samples the altitude of harvesting, both the maturity index (MI) and the altitude of harvesting data were taken from previous research (Reference).

### Statistical analysis

Dietary fiber data set of the fruit collection was analyzed by the Kolmogorov-Smirnov test for normality, one-way ANOVA, Box-plot representation, principal component analysis (PCA), and hierarchical clustering. Spearman’s correlation coefficients were calculated to evaluate potential dependencies among dietary fiber parameters and to determine whether those variables were related to harvesting altitude and MI. The mentioned statistical analyses were done using R-Studio software (2025.09.2+418), with the packages: ggplot2 [28], nortest [29], tidyr [30], dplyr [31], ggrepel [32], tibble [33], and ragg [34].

### Genotypic data, population structure, and genome-wide association analysis

The GWAS were performed using a previously generated single nucleotide polymorphism (SNP) dataset for *V. meridionale*, originally described by Narváez et al. (2026, accepted). Briefly, genotyping was conducted using a genotyping-by-sequencing (GBS) approach, and raw sequencing reads were processed to identify high-quality SNPs. Clean reads were mapped to the *V. corymbosum* cv. Draper v1.0 reference genome [35], using the set of the longest scaffolds (scaffolds 1 to 48) to represent the 48 chromosomes [36]. SNP markers with more than 30% missing data, those with a minor allele frequency (MAF) lower than 0.01, as well as INDELs, multiallelic, and monomorphic *loci* were excluded. In addition, genotypes presenting more than 70% missing data were excluded. Following quality filtering, 69 of the initial 118 phenotyped genotypes were retained, yielding 16,709 high-quality SNPs across the filtered genotypes for downstream analyses.

Population genetic structure, genome linkage disequilibrium (LD), and genome-wide SNP distribution were previously characterized using this SNP dataset, as described by Narváez et al. [10]. Population structure was inferred using admixture analysis implemented through the sNMF algorithm, and ancestry coefficients and LD estimates were directly incorporated into the association models to control for population stratification.

To identify significant quantitative trait loci (QTL) associations, a GWAS were performed independently for each phenotypic trait that exhibited significant variability among the 69 genotypes, as determined by ANOVA, using seven different association mapping models, as follows: General Linear Model (GLM) [37], Mixed Linear Model (MLM) [38], Multiple Loci Mixed Model (MLMM) [39], Compressed Mixed Linear Model (CMLM) [40], Fixed and Random Model Circulating Probability Unification (FarmCPU) [41], Bayesian-information and Linkage-disequilibrium Iteratively Nested Keyway (BLINK) [42], and Settlement of MLM Under Progressively Exclusive Relationship (SUPER) [43]. Significance of marker–trait associations was initially assessed using nominal *p*-values derived from the GWAS models. Multiple testing was accounted for by applying a false discovery rate (FDR) correction, and associations with FDR-adjusted *p-value* < 0.05 were retained as significant. Quantile–quantile plots were examined to evaluate model performance and control of Type I error [44]

### Candidate gene identification

Genomic positions of the identified QTLs were mapped and predicted functional annotation was performed to classify each variant as intergenic, intronic, coding, or regulatory. using the *V. corymbosum* cv. Draper v1.0 reference genome [35]. Candidate genes were identified within a LD window of ±161.782 kb surrounding each significant QTL, according to the LD decay previously reported for the genotype-population [10]. Gene annotation information was retrieved from the VacciniumGDB database (https://www.vaccinium.org/).

Functional annotation of candidate genes was performed using protein homology and curated biological information from the UniProt (https://www.uniprot.org/) and NCBI (https://www.ncbi.nlm.nih.gov/) databases. Gene Ontology (GO) analyses were conducted using the OmicsBox cloud platform (https://www.biobam.com/omicsbox/).

## Results

### Dietary fiber fractions across the fruit collection

Information on the SDF, IDF, TDF contents (in DW) as well as the SDF/IDF * 100 ratio is presented in S1 Table. While in the fruit collection the IDF content ranged from 19.9 ± 1.2 g/100 g DW (in G13) to 47.0 ± 2.0 g/100 g DW (in E02) (2.4-fold change), the SDF content ranged from 0.9 ± 0.2 g/100 g DW (in CP 23) and 0.9 ± 0.6 g/100 g DW (in CP40) to 6.1 ± 1.6 g/100 g DW (in G08) (6.5-fold change). Additionally, the TDF content ranged from 23.1 ± 2.0 g/100 g DW (in G03) to 49.6 ± 1.4 g/100 g DW (in E02). In all cases, the IDF fraction showed higher levels than the SDF fraction, with a SDF/IDF * 100 ratio ranging from 3.0 ± 1.8% (in SC22) to 24.9 ± 4.9% (in G08), highlighting a compositional profile dominated by the IDF. The coefficients of variation (CV) associated with biological replicates ranged from 0.6 to 25.7% for IDF, 3.5 to 97.2% for SDF, and 0.8 to 22.2% for TDF. In contrast, CV values obtained from technical replicates were lower, ranging from 0.34 to 10.4% for IDF, 1.3 to 24.9% for SDF, and 0.1 to 9.3% for TDF. Furthermore, The IDF and TDF values recorded in the control fruit sample (29.7 ± 0.5 and 30.6 ± 0.2 g/100 g DW, respectively) fall within the mid-range of the values detected across the fruit collection. In contrast, the SDF content in such control sample (1.0 ± 0.4 g/100 g DW) is at the lowest end of the observed range for the fruit collection. When analyzing data of the fruit collection together with the control sample, the ANOVA indicated that IDF, SDF, TDF, and the SDF/IDF * 100 ratio were statistically influenced by the genotype (F value < 0.0001). Considering a serving size of approximately 150 g (FW) and a recommended daily total dietary fiber intake of 30 g [9], the control ripe fruit sample, with a content of 3.3 ± 1.3 g TDF/100 g FW, would contribute only about 17% of the daily requirement. In contrast, several genotypes within the fruit collection showed a markedly higher nutritional contribution. Four genotypes supplied 90-100% of the recommended daily intake (SC16, 93%; G15, 97%; G21, 99%; and GA08, 100%). Furthermore, five genotypes exceeded this threshold, providing more than 100% (G18, 102%; G13, 108%; G24, 108%; GA24, 120%; and G08, 124%).

### Correlations among variables

No significant associations were found between dietary fiber fractions (IDF, SDF, TDF) or the SDF/IDF × 100 ratio and the altitude at which fruits were collected, suggesting that altitude did not substantially influence fiber composition under the conditions studied. In contrast, the MI was positively associated with IDF (r = 0.2655, *p* < 0.01), SDF (r = 0.2217, *p* < 0.05), and TDF (r = 0.3003, *p* < 0.001). A strong and significant positive correlation was found between IDF and TDF (r = 0.9622, *p* < 0.001), reflecting the predominance of the IDF over the SDF content across samples. Furthermore, SDF showed a positive, strong, and significant correlation with the SDF/IDF * 100 ratio (r = 0.9128, *p* < 0.001). A negative, strong, and significant correlation was found between IDF and the ratio SDF/IDF * 100 (r = -0.5661, *p* < 0.001). All these results are presented in S1 Fig.

### Principal component analysis (PCA) and cluster analysis

PCA revealed that the first two components accounted for 99.5% of the total inertia (Fig 1). Proximity of vectors confirm the relationship described in the precedent subheading between IDF and TDF as well as between SDF and SDF/IDF * 100 ratio. Clustering analysis yielded three groups with biological interest. Group I (blue) comprises 14 genotypes and was characterized by the greatest IDF and TDF contents, along with intermediate SDF content and SDF/IDF x 100 ratio. The elevated TDF values observed in genotypes E02 (49.6 ± 1.3 g/100 g DW), O02 (47.5 ± 0.7 g/100 g DW), B07 (47.5 ± 1.1 g/100 g DW), B06 (47.4 ± 5.6 g/100 g DW), and T02 (45.1 ± 1.5 g/100 g DW), identified as outliers in the box-plot analysis (S2 Fig), suggest a superior capacity for dietary fiber accumulation, which may be associated with genotype-dependent differences in cell wall composition. Group II (green) includes the majority of the genotypes (82 out of the 119 fruit material collected) and was characterized by intermediate IDF and TDF contents and the lowest SDF content and SDF/IDF × 100 ratio. In contrast, Group III (orange), comprising 23 genotypes, showed the lowest IDF and TDF contents but the greatest SDF content and SDF/IDF × 100 ratio. Within this group, genotypes G08 and GA24 exhibited the greatest SDF contents (6.1 ± 1.5 g/100 g DW and (6.1 ± 0.4 g/100 g DW) and the greatest SDF/IDF × 100 ratios (24.9 ± 4.9% and 24.0 ± 3.2%, respectively). These genotypes were identified as outliers in the Box-plot analysis (S2 Fig), indicating a distinct fiber composition profile dominated by soluble fractions. Taken together, Groups I and III represent contrasting dietary fiber profiles, with Group I characterized by a high TDF and IDF content, whereas Group III is distinguished by a higher proportion of soluble fiber.

**Fig 1.**
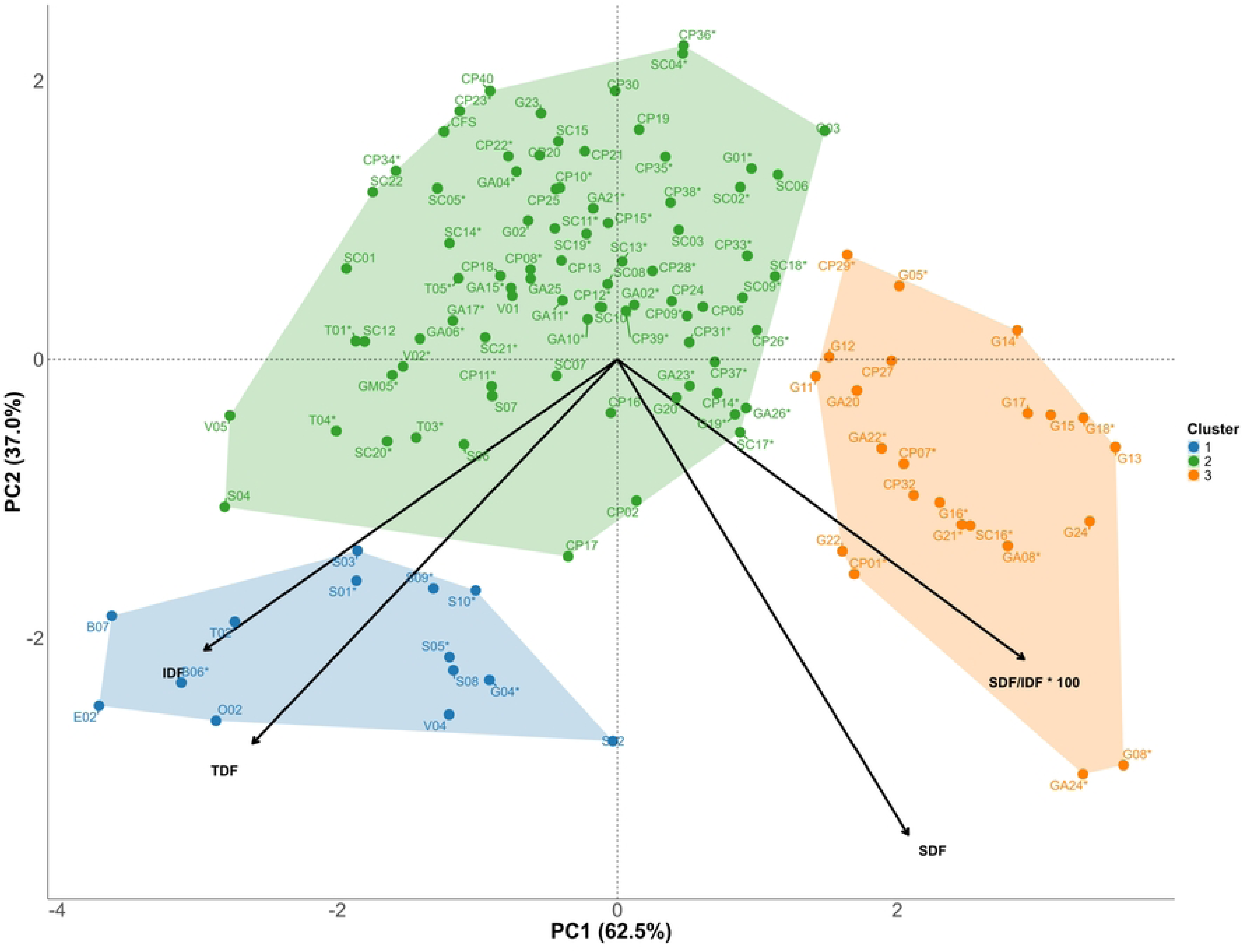
Principal component analysis (PCA) biplot with clustering based on insoluble dietary fiber (IDF), soluble dietary fiber (SDF), total dietary fiber (TDF), and the SDF/IDF × 100 ratio. Vectors representing each dietary fiber variable are shown. Three distinct clusters were identified. Group I (blue) comprises 14 genotypes and is characterized by the greatest IDF and TDF contents, along with intermediate SDF content and SDF/IDF x 100 ratio. Group II (green) includes 82 genotypes and is characterized by intermediate IDF and TDF contents and the lowest SDF content and SDF/IDF × 100 ratio. Group III (orange), comprising 23 genotypes, is characterized by the lowest IDF and TDF contents but the greatest SDF content and SDF/IDF × 100 ratio. Genotypes marked with an asterisk are those included in the genome-wide association study (GWAS).

### Genome-wide association analysis

A total of 24 QTLs were identified across the evaluated traits, showing levels of significance ranging from 3.39 × 10⁻¹⁰ to 2.71 × 10⁻⁷. Among these, the QTL Chr10:31193910 was detected in association with both TDF and IDF, exhibiting the highest statistical significance, whereas the QTL Chr41:26883013, associated with TDF, showed the lowest significant value (Table 1).

**Table 1.**
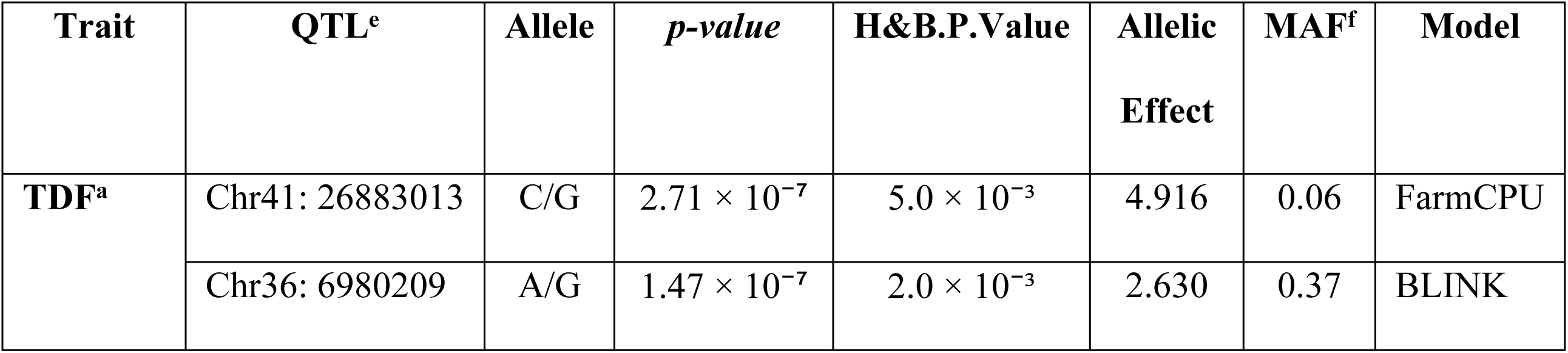
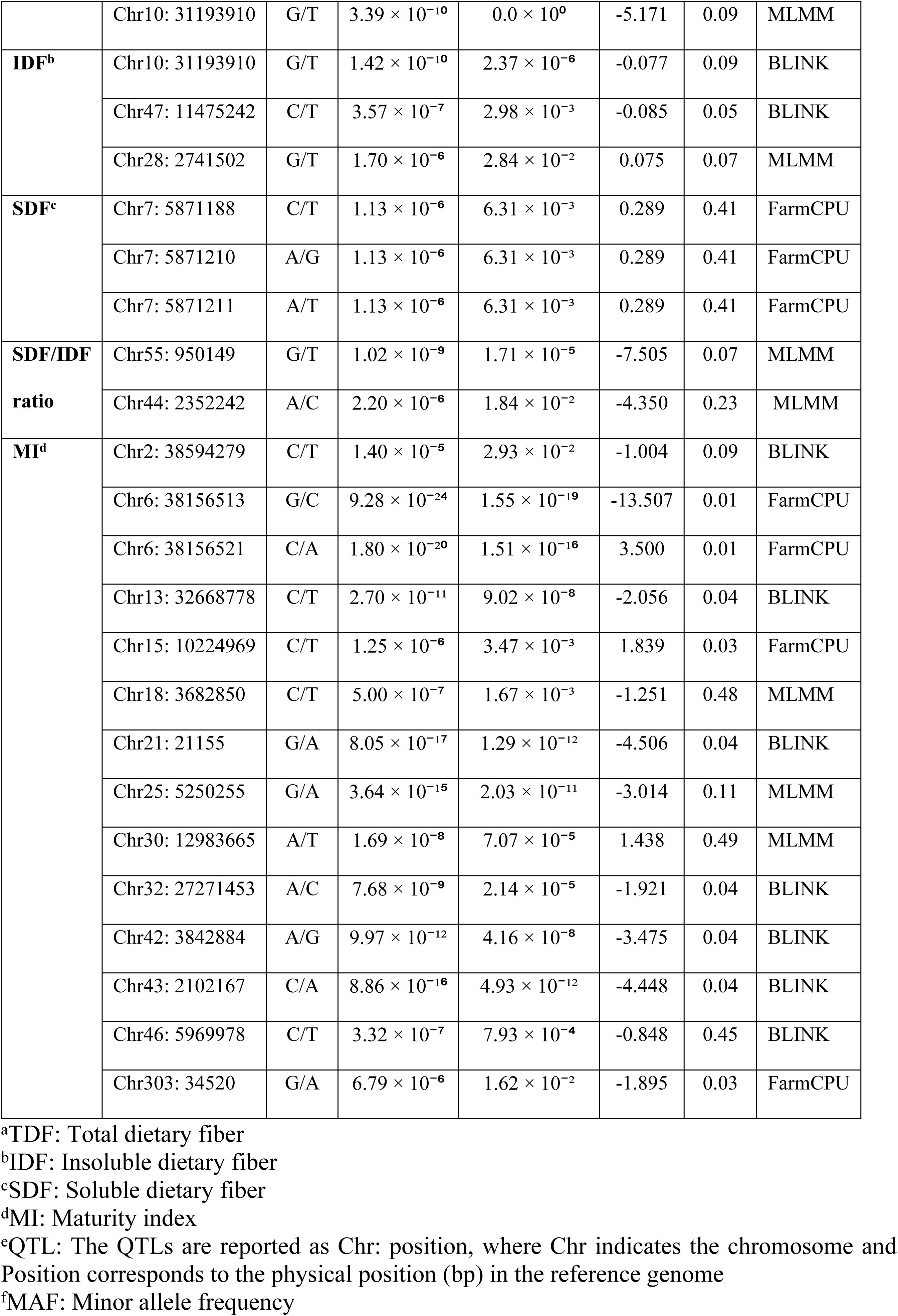
Statistical summary of QTLs significantly associated with dietary fiber-related traits identified by GWAS in *Vaccinium meridionale* populations.

Significant associations were detected using the MLMM, BLINK, and FarmCPU models after correction for population structure and multiple testing. The Q-Q plots indicated an adequate fit between observed and expected *p*-values for each analyzed trait, supporting the robustness of the applied association models (Figs 2 and 3). The Manhattan plots illustrate the genomic distribution of the significant QTLs across the genome (Figs 2 and 3).

**Fig 2.**
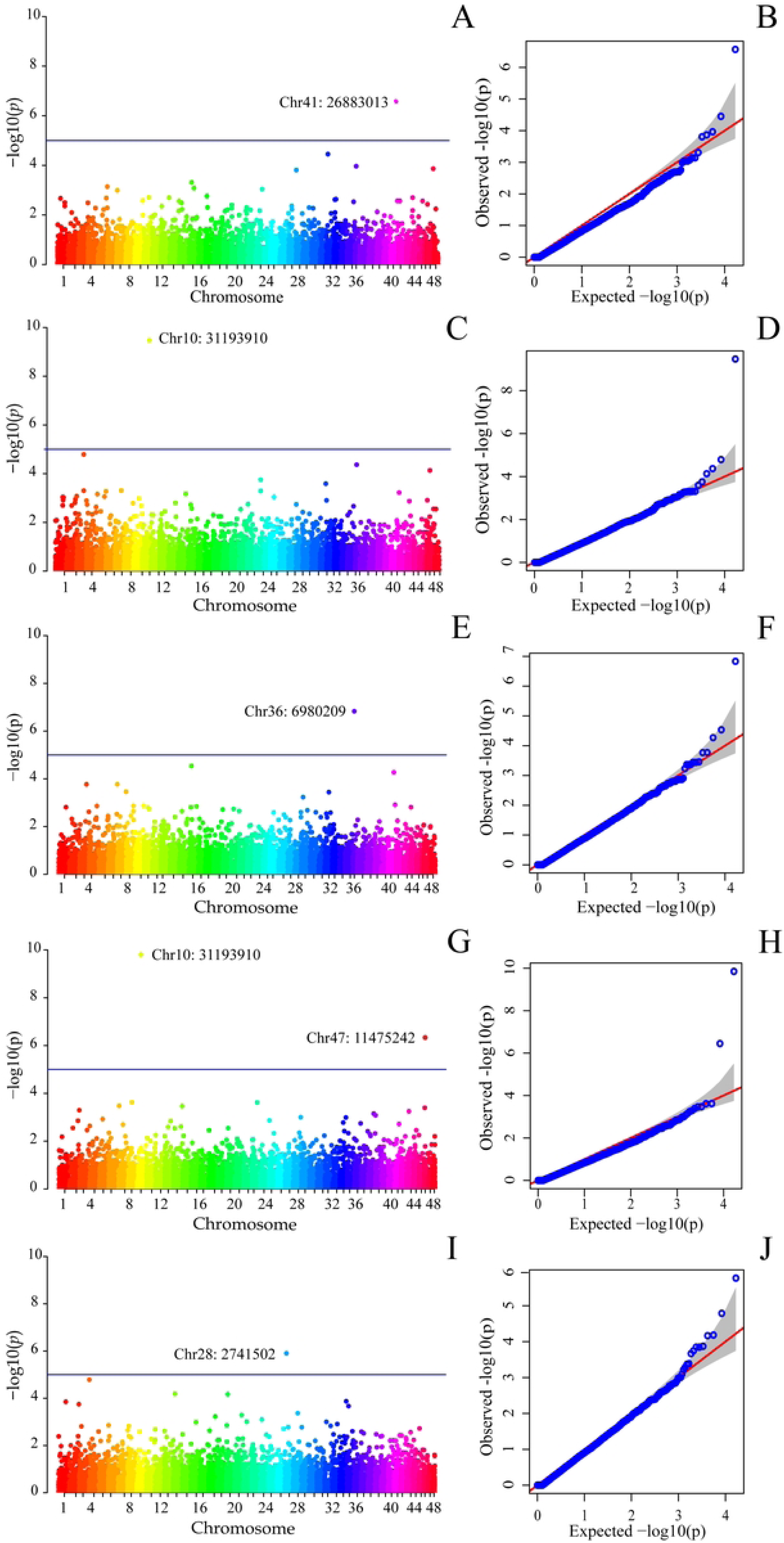
Manhattan and Q–Q plots of estimated –log10(*p-value*) for SNP associations with dietary fiber traits identified by GWAS. (A, B) Total dietary fiber (TDF) by the FarmCPU model. (C, D) TDF by the MLMM model. (E, F) TDF by the BLINK model. (G, H) Insoluble dietary fiber (IDF) by the BLINK model. (I, J) IDF by the MLMM model. The horizontal dashed line indicates the genome-wide significance threshold for phenotype–genotype associations.

**Fig 3.**
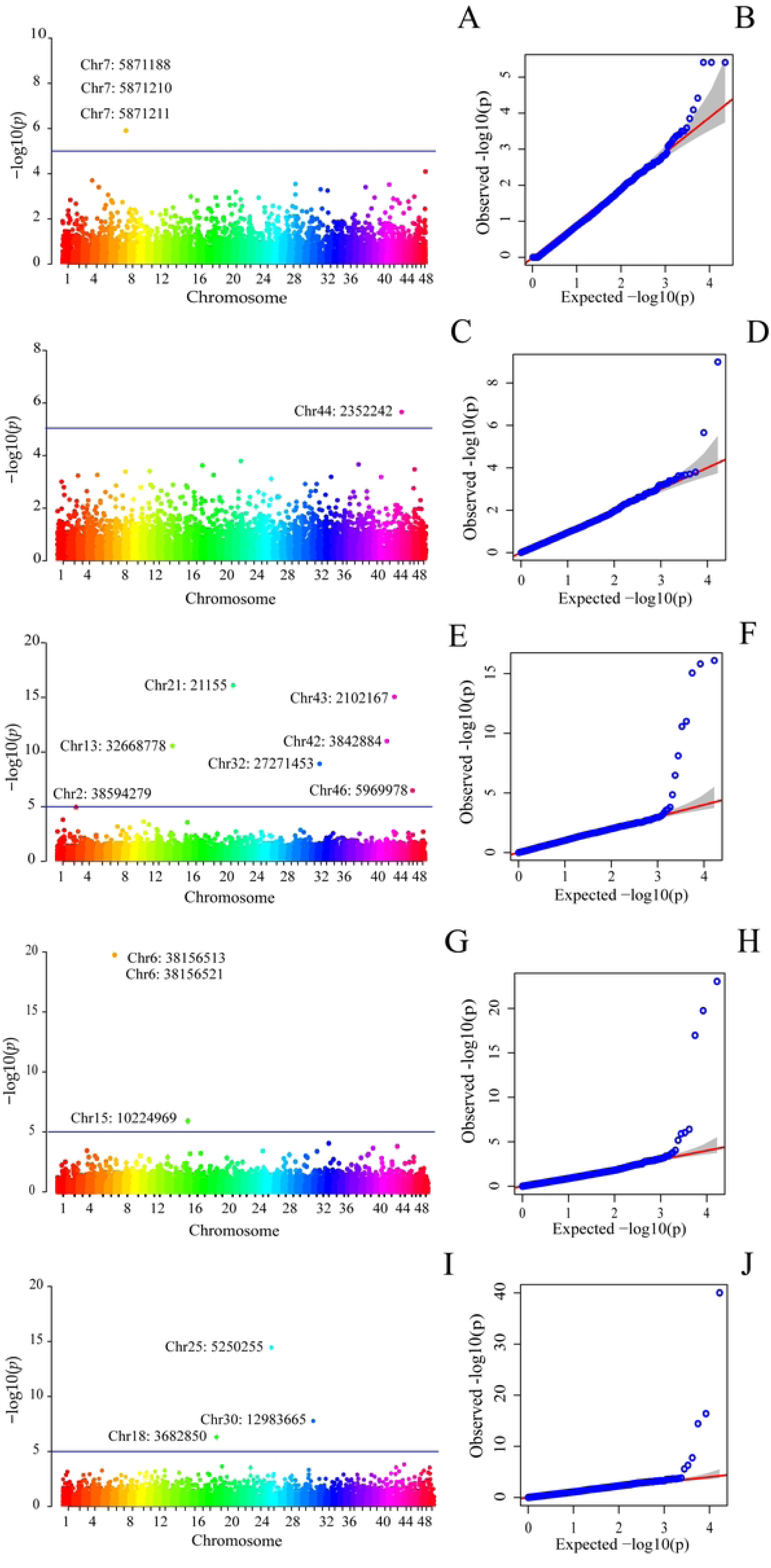
Manhattan and Q–Q plots of estimated –log10(*p-value*) for SNP associations with dietary fiber traits identified by GWAS. (A, B) Soluble dietary fiber (SDF) by the FarmCPU model. (C, D) SDF/IDF × 100 ratio by the MLMM model. (E, F) maturity index (MI) by the BLINK model. (G, H) MI by the FarmCPU model. (I, J) MI by the MLMM model. The horizontal dashed line indicates the genome-wide significance threshold for phenotype–genotype associations.

Regarding their genomic distribution, the QTLs were located in different functional regions, including nine QTLs in introns, nine within coding sequences (CDS), one in the 5′ UTR, three in the 3′ UTR, and two in intergenic regions. Among the QTLs located in coding regions, five nucleotide substitutions resulted in non-synonymous amino acid changes, while four corresponded to synonymous substitutions (Table 2).

**Table 2.**
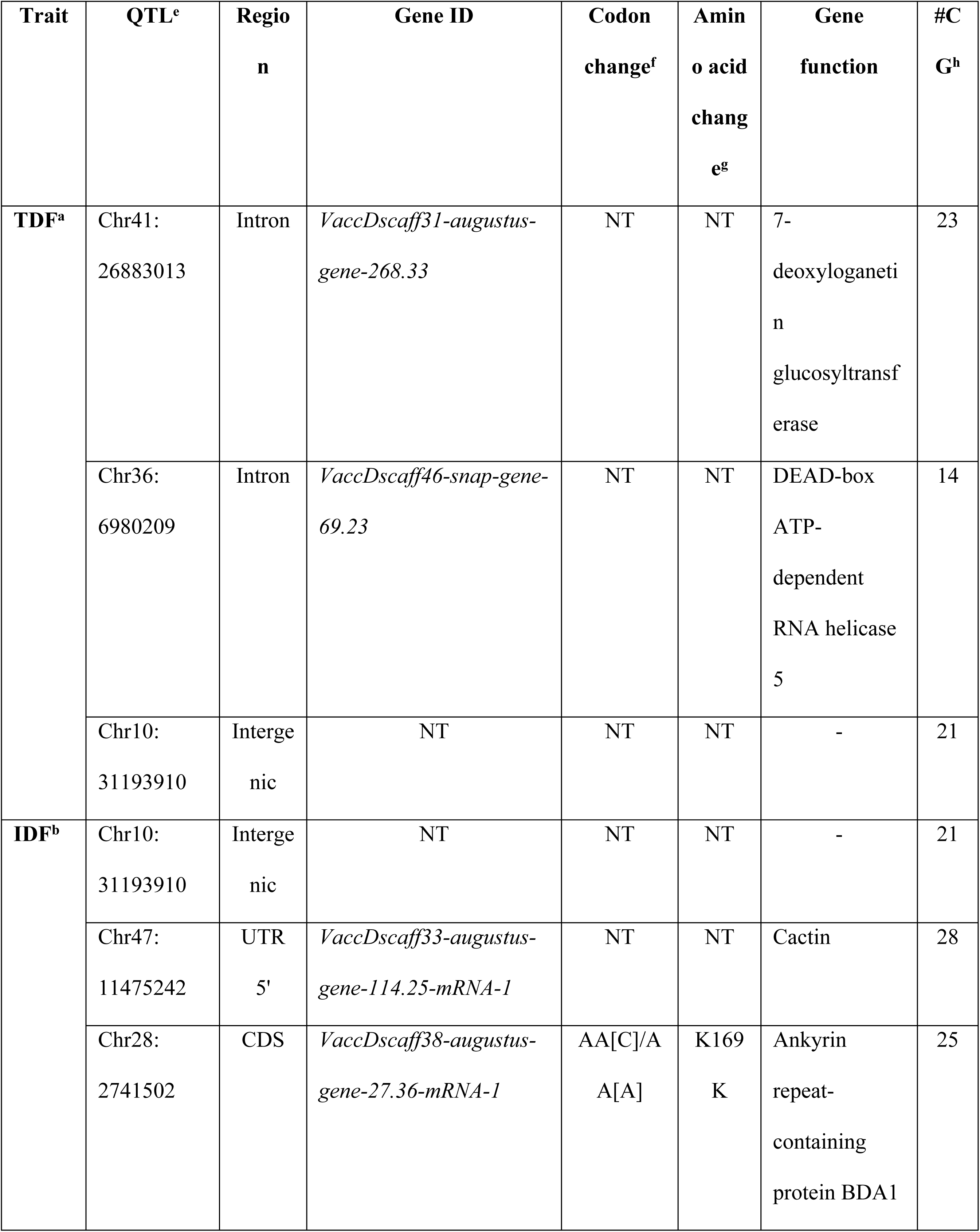
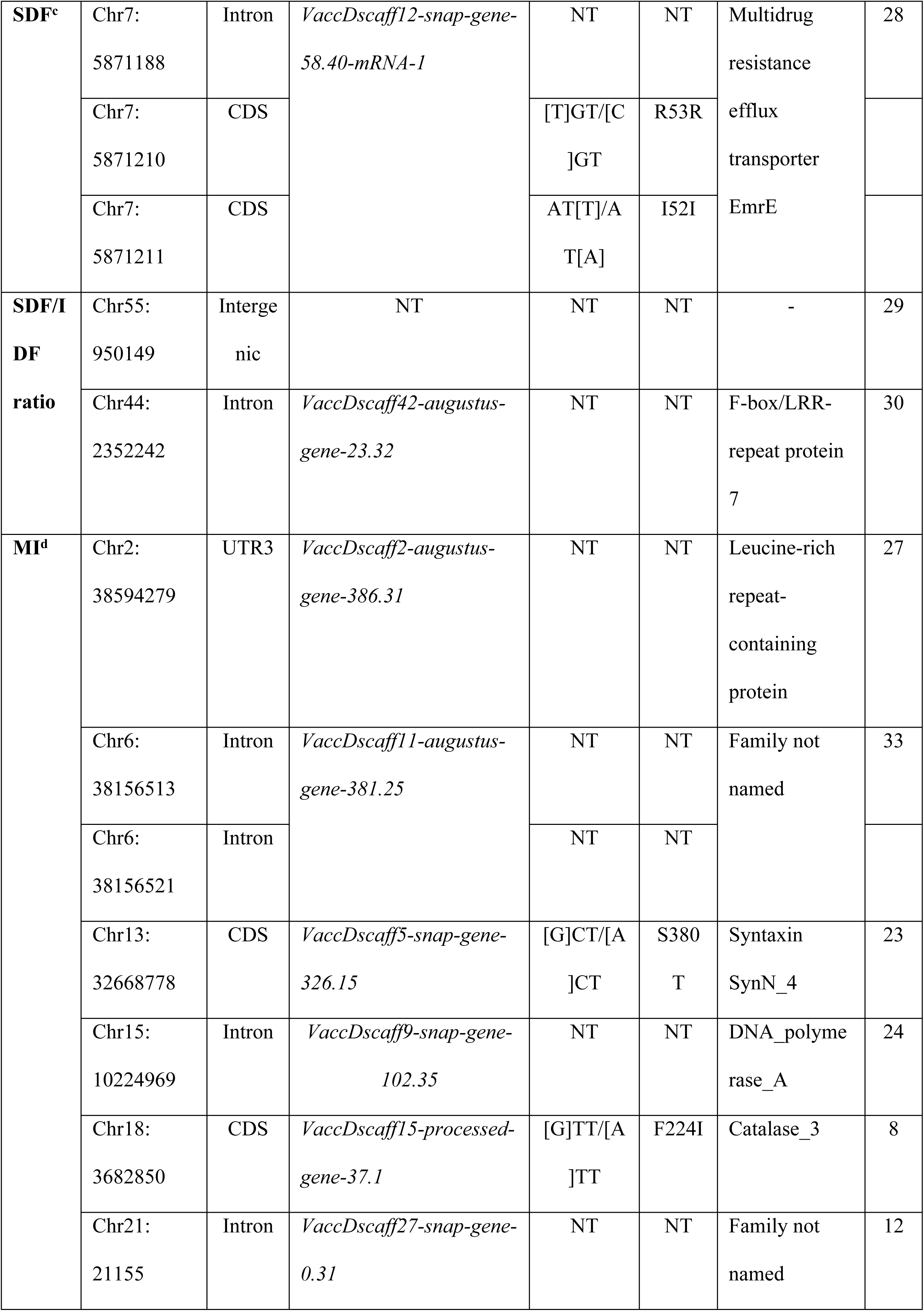
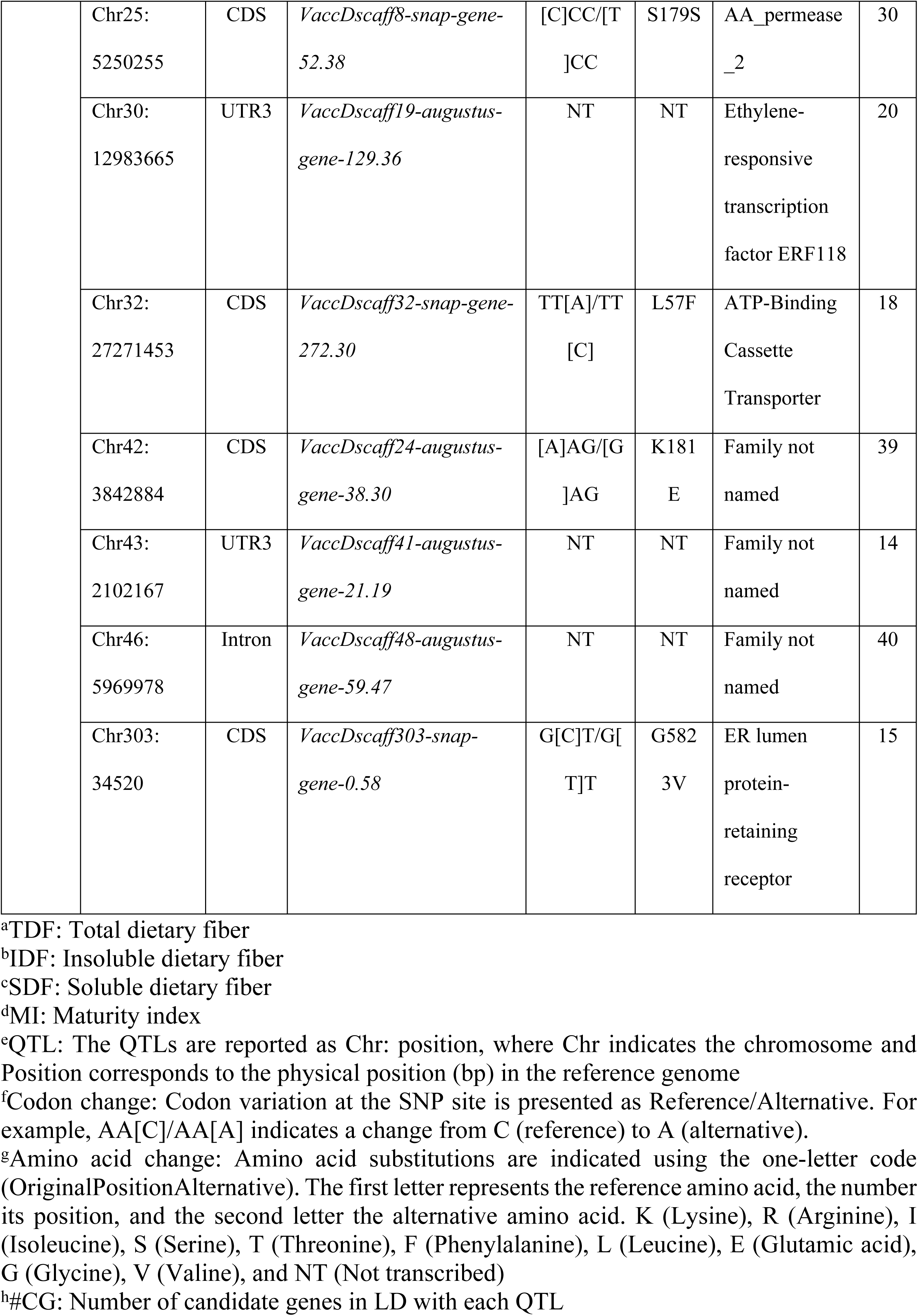
Functional annotation and predicted coding effects of QTL-associated candidate genes identified in *Vaccinium meridionale*.

Based on a linkage disequilibrium window of ±161.782 kb, a total of 501 candidate genes were identified across all traits. These genes were defined exclusively by their physical proximity to significant QTLs (S2 Table).

### Functional characterization of candidate genes based on Gene Ontology

Among the 501 analyzed genes, 380 were successfully annotated with Gene Ontology terms. Within the Cellular Component category, the most represented terms were intracellular anatomical structure (119 genes), membrane (108), organelle (100), and cytoplasm (76), while a smaller proportion of genes was associated with the endomembrane system (22). In the Molecular Function category, genes were predominantly involved in binding and enzymatic activities, with organic cyclic compound binding (83 sequences) and small molecule binding (79) being the most frequent, followed by transferase activity (54), hydrolase activity (36), catalytic activity acting on a protein (33), carbohydrate derivative binding (30), and oxidoreductase activity (20). Regarding Biological Process, the dominant term was metabolic process (118 sequences), accompanied by a substantial representation of regulation of cellular process (50), establishment of localization (35), cellular response to stimulus (33), response to stress (30), and cellular component organization or biogenesis (26), whereas transmembrane transport, signaling, signal transduction, and cell communication were each represented by 22 sequences (Fig 4).

**Fig 4.**
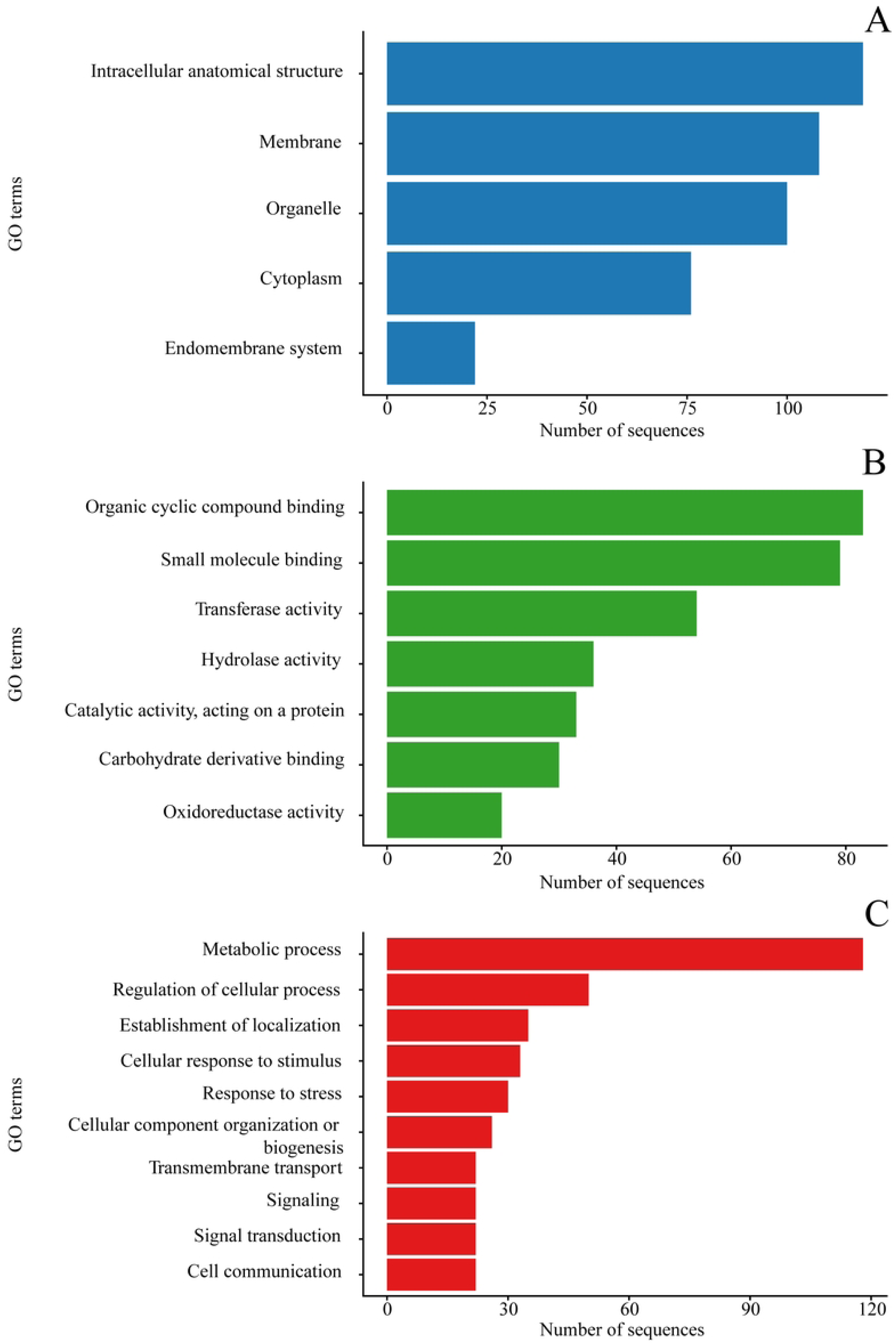
Gene Ontology (GO) classification of the annotated genes. Bar plots show the distribution of GO terms assigned to the 380 annotated genes across the three main GO categories: (A) Cellular Component, (B) Molecular Function, and (C) Biological Process. While the x-axis indicates the number of sequences associated with each term, the y-axis represents the GO terms within each category.

## Discussion

### Genotype and physiological maturity drive dietary fiber variability in *V. meridionale* fruits

The contents of IDF found in the fruit collection of *V. meridionale* are greater than the values found in six cultivars of *V. corymbosum* (blueberry, 10.6 -15.5 g/100 g DW) [45], and also greater than the one found in *Vaccinium myrtillus* (bilberry, 17.8 g/100 g DW) [46]. In contrast, the values reported of SDF not only in *V. myrtillus* (6.2 g/100 g DW) [46] but also in *V. corymbosum* (six cultivars, 2.4 -4.3 g/100 g DW) [45] are comparable with the values for this dietary component found in the *V. meridionale* fruit collection. Predominance of IDF towards SDF coincides with such findings in *V. corymbosum* [45] and *V. myrtillus* (Reference) and it seems to be a common feature in different fruits [47]. The lower variability observed between technical replicates compared to biological replicates supports the reproducibility of the analytical technique used. Therefore, the high dispersion obtained between genotypes, mainly for SDF content, is attributed to biological variability and not to methodological limitations. IDF is associated with a positive effect on bulking fecal material, improves gastrointestinal motility, can be fermented at a certain extent by gut microbiota, and is more protective than SDFs against some metabolic diseases such as Type 2 diabetes mellitus. In contrast, SDF reduces the diffusion and absorption rates of glucose and lipids and is closely related to improving gut microbiota due to its high fermentability which is negatively correlated with the incidence rate of many chronic diseases, including inflammatory bowel disease, colorectal cancer, obesity, and type 2 diabetes mellitus [48,49]. This divergence highlights complementary nutritional targets for genotype selection.

Moreover, the identified genotypes meeting daily dietary fiber requirements suggest that these materials would provide good candidates for breeding programs.

Although fruits were harvested at a similar external color, the MI exhibited considerable variability, suggesting heterogeneity in physiological maturity. A previous study analyzed ripe berries from a large set of *Vaccinium stamineum* genotypes harvested when pedicels changed from green to red, which was used as a visual indicator of maturation. However, despite the application of this criterion, considerable variability in physiological maturity was detected, as reflected by differences in pH, total soluble solids, and titratable acidity [50]. The authors concluded that both genotype and physiological maturity significantly affected these quality parameters. These findings are consistent with the variability observed in the present study and support the notion that visual harvest indices alone may not adequately represent physiological maturity in Vaccinium berries. Additionally, the MI may contribute to differences in the accumulation or structural modification of dietary fiber components, potentially through ripening associated cell wall polysaccharide remodeling [51].

### Multiple QTLs reveal polygenic and regulatory control of dietary fiber composition in *V. meridionale*

Genome-wide association analyses confirmed a complex and polygenic genetic architecture underlying dietary fiber–related traits in *V. meridionale*. Across TDF, IDF, SDF, the SDF/IDF ratio, and MI, multiple QTLs were identified across at least 15 chromosomes using complementary association models (BLINK, FarmCPU, and MLMM). These QTLs mapped to both coding and non-coding genomic regions, supporting a genetic model in which dietary fiber traits are shaped by a combination of structural variation in key enzymes and regulatory variation affecting gene expression [52]. This pattern is consistent with the quantitative nature of dietary fiber composition, which depends on coordinated cell wall biosynthesis, remodeling, and developmental regulation rather than on single major-effect loci [53].

### Glycosyltransferase (GT) gene families underpin fiber-related variation

A striking convergence between QTL location and functional annotation was observed for genes encoding glycosyltransferases across all fiber-related traits, supporting their central role in dietary fiber variation in *V. meridionale*. For TDF, the QTL Chr41:26883013 directly overlapped the glycosyltransferase gene *VaccDscaff31-augustus-gene-268.33*, annotated as a 7-deoxyloganetin glucosyltransferase and showing 56.31% sequence identity with its *Gardenia jasminoides* homolog, strongly suggesting this gene as a primary causal candidate underlying the QTL effect. This signal was further reinforced by the presence of five additional glycosyltransferase genes within the same LD block, spanning distances from 11.4 to 52.7 kb, including *VaccDscaff31-processed-gene-268.6* (11,444 bp), *VaccDscaff31-snap-gene-268.40* (16,617 bp), and *VaccDscaff31-augustus-gene-268.31* (40,394 bp), all annotated as 7-deoxyloganetin glucosyltransferases, as well as *VaccDscaff31-augustus-gene-268.38* (52,534 bp), encoding a UDP-glycosyltransferase 85A3, and *VaccDscaff31-processed-gene-268.14* (52,689 bp), encoding UDP-glycosyltransferase 85A5. The tight physical clustering and functional coherence of these genes indicate a glycosylation hotspot likely contributing to variation in fiber biosynthesis or cell wall-associated secondary metabolites.

For IDF, the QTL Chr47:11475242 mapped within a LD block containing *VaccDscaff33-snap-gene-113.46*, located 99.5 kb from the peak SNP and annotated as a probable glycosyltransferase homologous to *At5g03795* from *Arabidopsis thaliana*. Although this gene does not overlap the QTL peak, its proximity and annotation support a role in the biosynthesis and modification of hemicellulosic polysaccharides, tightly bound to the cell wall matrix, and cellulosic polysaccharides, which constitute the main structural components of IDF [54,55]. In addition, glycosyltransferase mediated glycosylation has been shown to regulate lignin deposition and secondary wall development in fruit tissues, as demonstrated for *PbUGT72AJ2* in pear (*Pyrus bretschneideri)*, where altered uridine diphosphate glycosyltransferase (UGT) activity modulates lignin accumulation [56].

Similarly, for the SDF/IDF ratio, the QTL Chr55:950149 was associated with *VaccDscaff55-augustus-gene-9.30* at 29.3 kb, encoding a xyloglucan endotransglucosylase/hydrolase. Xyloglucan-modifying enzymes regulate hemicellulose restructuring and wall extensibility, processes known to influence fiber solubility [57,58]. The physical proximity of this gene to the QTL therefore supports its candidacy as a contributor to variation in the relative proportions of soluble and insoluble dietary fiber.

Glycosyltransferase involvement extended to MI, as MI-associated QTLs colocalized with genes encoding enzymes involved in cell wall polysaccharide modification rather than pathways directly related to sugar or organic acid biosynthesis. Specifically, the QTL Chr18: 3682850 was associated with *VaccDscaff15-snap-gene-38.27* (14.5 kb), encoding UDP-glycosyltransferase 92A1; Chr32: 27271453 with *VaccDscaff32-augustus-gene-273.17* (85.2 kb), encoding a dolichyl-diphosphooligosaccharide–protein glycosyltransferase subunit 4; and Chr43: 2102167 with *VaccDscaff41-augustus-gene-22.14* (124.9 kb), encoding a probable xyloglucan glycosyltransferase 12. These associations reinforce the role of glycosylation-driven cell wall remodeling during fruit maturation, a process in which xyloglucan-modifying enzymes are central to the assembly and restructuring of hemicellulose networks within the primary cell wall. Consistent with this interpretation, Yuan et al. [59] identified xyloglucan glycosyltransferases as core components of the cellulose–hemicellulose matrix, directly contributing to wall extensibility and polysaccharide architecture. Moreover, the significant positive correlations observed between MI and IDF, SDF, and TDF indicate that MI in *V. meridionale* integrates coordinated changes in dietary fiber composition, reflecting structural modifications of the cell wall that are characteristic of fruit ripening rather than solely the balance between soluble sugars and organic acids.

The enrichment of glycosyltransferase genes within fiber-associated QTLs is supported by functional studies demonstrating that GT involved in the assembly and remodeling of wall polysaccharides strongly influence fiber solubility and physicochemical properties [45]. Moreover, genome-wide characterization of the UDP-glycosyltransferase family (UGT subfamily) in blueberry (*V. corymbosum*) identified 361 VcUGTs, with marked expansion of groups E and G driven by genome duplication events [60]. The co-localization of UGTs within LD windows of multiple fiber-related QTLs in *V. meridionale* is therefore consistent with the evolutionary and functional prominence of this family in blueberry.

Functional evidence further supports this link: silencing of *VcUGT160* (UGT85 subfamily) altered cytokinin homeostasis during blueberry fruit development [60]. Given the role of cytokinins in cell expansion and wall deposition, allelic variation in UGT genes near fiber-associated QTLs may influence dietary fiber traits through hormone-mediated developmental pathways. Collectively, the repeated association of specific glycosyltransferase families; UDP-glycosyltransferases, and xyloglucan endotransglucosylase/hydrolases with QTLs across multiple traits provides strong evidence that allelic variation in glycosylation pathways represents a major genetic determinant of dietary fiber composition in *V. meridionale*.

### Cell wall remodeling enzymes and polysaccharide turnover

The QTLs identified for dietary fiber traits in *V. meridionale* revealed a consistent enrichment of candidate genes involved in cell wall remodeling and polysaccharide turnover, highlighting the importance of post-synthetic regulation of wall architecture in shaping fiber composition. Dietary fiber composition in fruits is increasingly recognized as a dynamic trait governed not only by primary biosynthetic pathways but also by extensive remodeling of pectic [61] and hemicellulosic polymers during development and ripening, which directly affects polysaccharide interactions, extractability, and solubility [52]. Consequently, the fiber-associated loci identified here point out to coordinated enzymatic and regulatory mechanisms controlling wall restructuring, rather than isolated variation in biosynthetic capacity.

A key component of this regulatory framework involves genes controlling pectin modification, given the central role of pectin structure in cell wall mechanics and fruit texture [62]. The IDF-associated QTL Chr47: 11475242 was linked to *VaccDscaff33-processed-gene-116.2*, located 143.3 kb from the peak SNP and annotated as a probable pectin methylesterase. Although pectins are a major component of SDF, their physicochemical state is highly dynamic and can be altered by enzymatic modification, particularly through changes in the degree of methyl-esterification that influence solubility and interactions within the cell wall [63,64]. Pectin methylesterases regulate the demethylesterification of homogalacturonan, a process that reduces pectin solubility by promoting calcium-mediated cross-linking and enhanced interactions with cellulose and hemicelluloses [62,65]. Consequently, increased demethylesterification favors the formation of rigid pectate networks with reduced extractability, effectively shifting pectins toward the insoluble dietary fiber fraction [66,67]. Consistent with previous studies linking cell wall pectin to tissue mechanical properties, and intercellular adhesion and firmness in fruits [68,69]. The proximity of this pectin methylesterase gene to the IDF-associated QTL supports its role in modulating insoluble fiber accumulation through coordinated regulation of cell wall architecture rather than direct polymer identity.

In addition to enzymatic candidates, several QTL regions harbored regulatory genes implicated in signaling pathways that coordinate cell wall remodeling with developmental cues. Notably, receptor-like kinases (RLKs) were associated with the IDF QTL Chr28:2741502, including *VaccDscaff38-snap-gene-27.46* at 29.7 kb, and with the MI QTL Chr25:5250255 through *VaccDscaff8-snap-gene-54.38* at 152.3 kb. RLKs form a large family of plasma membrane localized receptors that perceive extracellular and developmental signals and transduce them to intracellular signaling networks, thereby influencing cell expansion, wall integrity maintenance, and polysaccharide remodeling [70,71]. Functional studies have shown that specific RLK subfamilies, such as wall-associated kinases (WAKs) and malectin-like receptor kinases (CrRLK1Ls), directly interact with cell wall components, including pectin, and trigger downstream pathways that regulate cell wall dynamics and development [72,73]. For example, WAKs can bind pectin fragments resulting from wall modification and initiate signaling that alters expression of cell wall remodeling genes, while CrRLK1Ls such as FERONIA mediate growth regulation in response to cell wall status [72]. The co-localization of these receptor kinases with fiber-related QTLs in *V. meridionale* indicates that allelic variation in RLK-mediated signaling could underlie differential regulation of wall remodeling during fruit development and maturation, integrating environmental and developmental signals with changes in wall architecture.

Taken together, our results provide the first comprehensive insight into the polygenic basis of dietary fiber variation in *V. meridionale*, revealing a complex and promising genetic framework that opens new avenues for molecular and precision breeding strategies in agraz. Nonetheless, expanding population size and incorporating rigorous functional validation approaches will be essential to confirm causality and to fully establish the biological relevance of the identified candidate genes.

## Conclusion

This study presents an extensive and underexplored diversity in dietary fiber composition within *Vaccinium meridionale*, identifying this species as a promising source of dietary fiber within the Vaccinium genus. The identification of genotypes capable of meeting or exceeding recommended daily fiber intake, highlights the outstanding functional potential of this species. Moreover, the genetic architecture elucidated in this study indicates that cell wall glycosylation, remodeling processes, and putative regulatory pathways are major determinants of dietary fiber composition in *V. meridionale*. The consistent co-localization of fiber-related QTLs with genes encoding glycosyltransferases, pectin methylesterase enzymes, and xyloglucan-modifying enzymes provides promising targets for breeding strategies aimed at modulating both total fiber content and the balance between soluble and insoluble fractions. Given that dietary fiber strongly influences fruit texture, firmness, and extractability, allelic variation at these loci is expected to affect not only nutritional quality but also key fruit quality attributes relevant for fresh consumption and processing. In particular, genes involved in pectin and hemicellulose remodeling represent leverage points for fine-tuning fiber solubility, which are directly linked to recognized health benefits. Together, these results open future possibilities for developing a molecular framework to guide the breeding of Vaccinium varieties with improved dietary fiber profiles, with potential to integrate nutritional value and desirable fruit quality traits through marker-assisted and genomics-informed selection.

## Acknowledgments

The authors would like to thank the peasants in the various municipalities where the collection was carried out for their guidance and for serving as guides for the scientific team. We also thank the farmers who provided us with commercial materials.

## Supporting information

**S1 Table. Phenotypic dataset** of dietary fiber measurements in the *Vaccinium meridionale* population.

**S2 Table**: **List of candidate genes in linkage disequilibrium (LD)** surrounding significantly associated QTLs.

**S1 Fig**: **Spearmańs correlation coefficients** including IDF: Insoluble dietary fiber; SDF: Soluble dietary fiber; TDF: Total dietary fiber; SDF/IDF x 100 ratio, the altitude where each genotype was sampled and the maturity index. * Denotates *p* < 0.05; ** denotates *p* < 0.01; *** denotates *p* < 0.001;

**S2 Fig**: **Box-plot displayed for each of the measured response variables in the fruit collection**, including the control commercial fruit. IDF: Insoluble dietary fiber; SDF: Soluble dietary fiber; TDF: Total dietary fiber; and SDF/IDF x 100 ratio.

## Financial Disclosure Statement

This work was funded by the Ministry of Science, Technology, and Innovation of Colombia (MinCiencias) through the Sistema General de Regalías (SGR) under the project entitled “Aprovechamiento de la biodiversidad en agraz y papa para el desarrollo de cultivos promisorios en el departamento de Santander” (Project No. BPIN 2020000100075-2021), corresponding to the call “Convocatoria para la conformación de un listado de propuestas de proyectos elegibles para el fortalecimiento de capacidades institucionales y de investigación de las instituciones de educación superior públicas.”

The funders had no role in study design, data collection and analysis, decision to publish, or preparation of the manuscript.

## Data Availability Statement

The original contributions presented in this study are included in the article/Supplementary Materials. Further inquiries can be directed to the corresponding author.

## Competing interests

The authors declare no conflicts of interest.

## Author Contributions

Conceptualization: CENC, JCSS, TMV; Methodology: GPVA, ACGC; Software: GPVA, ACGC; Validation: CENC, JCSS; Formal analysis: GPVA, ACGC; Investigation: CENC, JCSS, GPVA, ACGC; Resources: TMV; Data curation: GPVA, ACGC, CENC; Writing – original draft preparation: GPVA; Writing – review & editing: CENC, JCSS, TMV; Supervision: CENC, JCSS, TMV; Project administration: CENC, TMV; Funding acquisition: TMV.All authors have read and agreed to the published version of the manuscript.

## Notes

### Competing Interest Statement

The authors have declared no competing interest.

## References

1. Montanari S, Thomson S, Cordiner S, Günther CS, Miller P, Deng CH, et al. High-density linkage map construction in an autotetraploid blueberry population and detection of quantitative trait loci for anthocyanin content. Front Plant Sci. 2022;13. doi:10.3389/fpls.2022.965397

2. Miller K, Feucht W, Schmid M. Bioactive Compounds of Strawberry and Blueberry and Their Potential Health Effects Based on Human Intervention Studies: A Brief Overview. Nutrients. 2019;11: 1510. doi:10.3390/nu11071510

3. Ligarreto-Moreno GA, Torres-Aponte WS, Ariza-Castillo A. Propagation of the neotropical fruit Vaccinium meridionale Swartz by air layering Propagación del frutal neotropical Vaccinium meridionale Swartz por acodos aéreos. 2013.

4. Garzón GA, Soto CY, López-R M, Riedl KM, Browmiller CR, Howard L. Phenolic profile, in vitro antimicrobial activity and antioxidant capacity of Vaccinium meridionale swartz pomace. Heliyon. 2020;6. doi:10.1016/j.heliyon.2020.e03845

5. Garzón GA, Soto CY, López-R M, Riedl KM, Browmiller CR, Howard L. Phenolic profile, in vitro antimicrobial activity and antioxidant capacity of Vaccinium meridionale swartz pomace. Heliyon. 2020;6. doi:10.1016/j.heliyon.2020.e03845

6. González M, Samudio I, Sequeda-Castañeda LG, Celis C, Iglesias J, Morales L. Cytotoxic and antioxidant capacity of extracts from Vaccinium meridionale Swartz (Ericaceae) in transformed leukemic cell lines. J Appl Pharm Sci. 2017;7: 24–30. doi:10.7324/JAPS.2017.70305

7. Arango-Varela SS, Luzardo-Ocampo I, Maldonado-Celis ME, Campos-Vega R. Andean berry (Vaccinium meridionale Swartz) juice in combination with Aspirin modulated anti-inflammatory markers on LPS-stimulated RAW 264.7 macrophages. Food Research International. 2020;137. doi:10.1016/j.foodres.2020.109541

8. Osorio M, Posada L, Martínez E, Estrada V, Quintana G, Maldonado ME, et al. Bacterial nanocellulose spheres coated with meta acrylic copolymer: Vaccinium meridionale swartz extract delivery for colorectal cancer chemoprevention. Food Hydrocoll. 2024;147. doi:10.1016/j.foodhyd.2023.109310

9. Veronese N, Gianfredi V, Solmi M, Barbagallo M, Dominguez LJ, Mandalà C, et al. The impact of dietary fiber consumption on human health: An umbrella review of evidence from 17,155,277 individuals. Clinical Nutrition. 2025;51: 325–333. doi:10.1016/j.clnu.2025.06.021

10. Narváez-Cuenca CE, Guevara-Correa AC, Velasco Anacona GP, Sepúlveda J, Soto Sedano JC, Mosquera-Vásquez T. New anthocyanin and non-anthocyanin phenolic compounds in wild bilberry (Vaccinium meridionale Swartz) and their genetic architecture. ACS Agricultural Science & Technology. In press.

11. Zhang X, Lu M, Ludlow RA, Ma W, An H. Transcriptome analysis reveals candidate genes for dietary fiber metabolism in Rosa roxburghii fruit grown under different light intensities. Hortic Environ Biotechnol. 2021;62: 751–764. doi:10.1007/s13580-021-00359-6

12. Du X, Pérez-Boada M, Fernández C, Rencoret J, del Río JC, Jiménez-Barbero J, et al. Analysis of lignin–carbohydrate and lignin–lignin linkages after hydrolase treatment of xylan–lignin, glucomannan–lignin and glucan–lignin complexes from spruce wood. Planta. 2014. doi:10.1007/s00425-014-2037-y

13. Mudgil D. The Interaction Between Insoluble and Soluble Fiber. Dietary Fiber for the Prevention of Cardiovascular Disease. Elsevier; 2017. pp. 35–59. doi:10.1016/B978-0-12-805130-6.00003-3

14. Schneider R, Hanak T, Persson S, Voigt CA. Cellulose and callose synthesis and organization in focus, what’s new? Current Opinion in Plant Biology. Elsevier Ltd; 2016. pp. 9–16. doi:10.1016/j.pbi.2016.07.007

15. Pauly M, Gille S, Liu L, Mansoori N, de Souza A, Schultink A, et al. Hemicellulose biosynthesis. Planta. 2013;238: 627–642. doi:10.1007/s00425-013-1921-1

16. Anders N, Wilson LFL, Sorieul M, Nikolovski N, Dupree P. β-1,4-Xylan backbone synthesis in higher plants: How complex can it be? Front Plant Sci. 2023;13. doi:10.3389/fpls.2022.1076298

17. Han Y, Zhu Q, Zhang Z, Meng K, Hou Y, Ban Q, et al. Analysis of Xyloglucan Endotransglycosylase/Hydrolase (XTH) Genes and Diverse Roles of Isoenzymes during Persimmon Fruit Development and Postharvest Softening. PLoS One. 2015;10: e0123668. doi:10.1371/journal.pone.0123668

18. Rohman A, Dijkstra BW, Puspaningsih NNT. β-Xylosidases: Structural Diversity, Catalytic Mechanism, and Inhibition by Monosaccharides. Int J Mol Sci. 2019;20: 5524. doi:10.3390/ijms20225524

19. Lagaert S, Pollet A, Courtin CM, Volckaert G. β-Xylosidases and α-l-arabinofuranosidases: Accessory enzymes for arabinoxylan degradation. Biotechnol Adv. 2014;32: 316–332. doi:10.1016/j.biotechadv.2013.11.005

20. Lu M, Ma W-T, Liu Y-Q, An H-M, Ludlow RA. Transcriptome Analysis Reveals Candidate Lignin-Related Genes and Transcription Factors in Rosa roxburghii During Fruit Ripening. Plant Mol Biol Report. 2020;38: 331–342. doi:10.1007/s11105-020-01193-3

21. Liu Y, Wang Y, Pei J, Li Y, Sun H. Genome-wide identification and characterization of COMT gene family during the development of blueberry fruit. BMC Plant Biol. 2021;21. doi:10.1186/s12870-020-02767-9

22. Ropartz D, Ralet M-C. Pectin Structure. Pectin: Technological and Physiological Properties. Cham: Springer International Publishing; 2020. pp. 17–36. doi:10.1007/978-3-030-53421-9_2

23. Lerouge P, Carlier M, Mollet J-C, Lehner A. The cell wall pectic rhamnogalacturonan II, an enigma in plant glycobiology. 2021. pp. 553–571. doi:10.1039/9781839164538-00553

24. Sterling JD, Atmodjo MA, Inwood SE, Kumar Kolli VS, Quigley HF, Hahn MG, et al. Functional identification of an *Arabidopsis* pectin biosynthetic homogalacturonan galacturonosyltransferase. Proceedings of the National Academy of Sciences. 2006;103: 5236–5241. doi:10.1073/pnas.0600120103

25. Wu H-C, Bulgakov VP, Jinn T-L. Pectin Methylesterases: Cell Wall Remodeling Proteins Are Required for Plant Response to Heat Stress. Front Plant Sci. 2018;9. doi:10.3389/fpls.2018.01612

26. Sepúlveda J, Rondón González F, Soto Sedano JC, Velasco GP, Mosquera T, Delgado MC, et al. SNP Analysis Reveals Novel Insights into the Genetic Diversity of Colombian Vaccinium meridionale. Genes (Basel). 2025;16. doi:10.3390/genes16060675

27. AOAC. Association of official analytical. Official methods of analysis (16th ed.). Gaithersburg,; 1995.

28. Wickham H. ggplot2. New York, NY: Springer New York; 2009. doi:10.1007/978-0-387-98141-3

29. Nortest: Gross J, & LU. nortest: Tests for normality. 2015.

30. Wickham H, VD, & GM. Tidyr: Tidy messy data (R package version 1.3.1). 2024.

31. Wickham H, FR, HL, MK, & VD. dplyr: A grammar of data manipulation. 2023.

32. Slowikowski K. ggrepel: Automatically position non-overlapping text labels with ‘ggplot2’ 2024.

33. Tibble: Müller K, & WH. tibble: Simple data frames. 2023.

34. Ragg: Pedersen TL, & SM. ragg: Graphic devices based on AGG. 2024.

35. Colle M, Leisner CP, Wai CM, Ou S, Bird KA, Wang J, et al. Haplotype-phased genome and evolution of phytonutrient pathways of tetraploid blueberry. Gigascience. 2019;8. doi:10.1093/gigascience/giz012

36. Nagasaka K, Nishiyama S, Fujikawa M, Yamane H, Shirasawa K, Babiker E, et al. Genome-Wide Identification of Loci Associated With Phenology-Related Traits and Their Adaptive Variations in a Highbush Blueberry Collection. Front Plant Sci. 2022;12. doi:10.3389/fpls.2021.793679

37. Price AL, Patterson NJ, Plenge RM, Weinblatt ME, Shadick NA, Reich D. Principal components analysis corrects for stratification in genome-wide association studies. Nat Genet. 2006;38: 904–909. doi:10.1038/ng1847

38. Yu J, Pressoir G, Briggs WH, Vroh Bi I, Yamasaki M, Doebley JF, et al. A unified mixed-model method for association mapping that accounts for multiple levels of relatedness. Nat Genet. 2006;38: 203–208. doi:10.1038/ng1702

39. Segura V, Vilhjálmsson BJ, Platt A, Korte A, Seren Ü, Long Q, et al. An efficient multi-locus mixed-model approach for genome-wide association studies in structured populations. Nat Genet. 2012;44: 825–830. doi:10.1038/ng.2314

40. Zhang Z, Ersoz E, Lai C-Q, Todhunter RJ, Tiwari HK, Gore MA, et al. Mixed linear model approach adapted for genome-wide association studies. Nat Genet. 2010;42: 355–360. doi:10.1038/ng.546

41. Liu X, Huang M, Fan B, Buckler ES, Zhang Z. Iterative Usage of Fixed and Random Effect Models for Powerful and Efficient Genome-Wide Association Studies. PLoS Genet. 2016;12: e1005767. doi:10.1371/journal.pgen.1005767

42. Huang M, Liu X, Zhou Y, Summers RM, Zhang Z. BLINK: a package for the next level of genome-wide association studies with both individuals and markers in the millions. Gigascience. 2019;8. doi:10.1093/gigascience/giy154

43. Wang Q, Tian F, Pan Y, Buckler ES, Zhang Z. A SUPER Powerful Method for Genome Wide Association Study. PLoS One. 2014;9: e107684. doi:10.1371/journal.pone.0107684

44. Turner SD. qqman: an R package for visualizing GWAS results using Q-Q and manhattan plots. 2014. doi:10.1101/005165

45. Chevalier LM, Rioux L, Angers P, Turgeon SL. Low-Temperature Blanching as a Tool to Modulate the Structure of Pectin in Blueberry Purees. J Food Sci. 2017;82: 2070–2077. doi:10.1111/1750-3841.13826

46. Aura AM, Holopainen-Mantila U, Sibakov J, Kössö T, Mokkila M, Kaisa P. Bilberry and bilberry press cake as sources of dietary fibre. Food Nutr Res. 2015;59. doi:10.3402/fnr.v59.28367

47. Li BW, Andrews KW, Pehrsson PR. Individual Sugars, Soluble, and Insoluble Dietary Fiber Contents of 70 High Consumption Foods. Journal of Food Composition and Analysis. 2002;15: 715–723. doi:10.1006/jfca.2002.1096

48. Guan Z-W, Yu E-Z, Feng Q. Soluble Dietary Fiber, One of the Most Important Nutrients for the Gut Microbiota. Molecules. 2021;26: 6802. doi:10.3390/molecules26226802

49. Li M, Ma S. A review of healthy role of dietary fiber in modulating chronic diseases. Food Research International. 2024;191: 114682. doi:10.1016/j.foodres.2024.114682

50. Matsumoto GO, Garcia A, Benevenuto J, Munoz PR. High-density linkage map and QTL analyses for fruit quality traits in the wild blueberry relative *Vaccinium stamineum*. G3: Genes, Genomes, Genetics. 2026;16. doi:10.1093/g3journal/jkaf263

51. Deng H, Wang X, Wang Y, Xiang Y, Chen M, Zhang H, et al. Dynamic Changes in Cell Wall Polysaccharides during Fruit Development and Ripening of Two Contrasting Loquat Cultivars and Associated Molecular Mechanisms. Foods. 2023;12: 309. doi:10.3390/foods12020309

52. Zhang X, Lu M, Ludlow RA, Ma W, An H. Transcriptome analysis reveals candidate genes for dietary fiber metabolism in Rosa roxburghii fruit grown under different light intensities. Hortic Environ Biotechnol. 2021;62: 751–764. doi:10.1007/s13580-021-00359-6

53. Persson S, Wei H, Milne J, Page GP, Somerville CR. Identification of genes required for cellulose synthesis by regression analysis of public microarray data sets. Proceedings of the National Academy of Sciences. 2005;102: 8633–8638. doi:10.1073/pnas.0503392102

54. Fangel JU, Petersen BL, Jensen NB, Willats WGT, Bacic A, Egelund J. A putative Arabidopsis thaliana glycosyltransferase, At4g01220, which is closely related to three plant cell wall-specific xylosyltransferases, is differentially expressed spatially and temporally. Plant Science. 2011;180: 470–479. doi:10.1016/j.plantsci.2010.11.002

55. Feng Y, Jin Q, Liu X, Lin T, Johnson A, Huang H. Advances in understanding dietary fiber: Classification, structural characterization, modification, and gut microbiome interactions. Compr Rev Food Sci Food Saf. 2025;24. doi:10.1111/1541-4337.70092

56. Wang H, Feng X, Zhang Y, Wei D, Zhang Y, Jin Q, et al. PbUGT72AJ2-Mediated Glycosylation Plays an Important Role in Lignin Formation and Stone Cell Development in Pears (Pyrus bretschneideri). Int J Mol Sci. 2022;23: 7893. doi:10.3390/ijms23147893

57. Stratilová B, Kozmon S, Stratilová E, Hrmova M. Plant Xyloglucan Xyloglucosyl Transferases and the Cell Wall Structure: Subtle but Significant. Molecules. 2020;25: 5619. doi:10.3390/molecules25235619

58. Vain T, Crowell EF, Timpano H, Biot E, Desprez T, Mansoori N, et al. The Cellulase KORRIGAN Is Part of the Cellulose Synthase Complex. Plant Physiol. 2014;165: 1521–1532. doi:10.1104/pp.114.241216

59. Yuan W, Yao F, Liu Y, Xiao H, Sun S, Jiang C, et al. Identification of the xyloglucan endotransglycosylase/hydrolase genes and the role of *PagXTH12* in drought resistance in poplar. Forestry Research. 2024;4: 0–0. doi:10.48130/forres-0024-0036

60. Wang Y, Liu X, Zhao T, Li J, Chen L, Li Y, et al. Genome-wide characterization of the UDP-glycosyltransferases (UGT) family and functional analysis of VcUGT160 involved in dihydrozeatin glycosylation during blueberry fruits development. BMC Genomics. 2025;26. doi:10.1186/s12864-025-12267-5

61. Di Matteo A, Giovane A, Raiola A, Camardella L, Bonivento D, De Lorenzo G, et al. Structural Basis for the Interaction between Pectin Methylesterase and a Specific Inhibitor Protein. Plant Cell. 2005;17: 849–858. doi:10.1105/tpc.104.028886

62. Huang W, Shi Y, Yan H, Wang H, Wu D, Grierson D, et al. The calcium-mediated homogalacturonan pectin complexation in cell walls contributes the firmness increase in loquat fruit during postharvest storage. J Adv Res. 2023;49: 47–62. doi:10.1016/j.jare.2022.09.009

63. Yüksel E, Kort R, Voragen AGJ. Structure and degradation dynamics of dietary pectin. Crit Rev Food Sci Nutr. 2025;65: 6249–6268. doi:10.1080/10408398.2024.2437573

64. Jolie RP, Duvetter T, Van Loey AM, Hendrickx ME. Pectin methylesterase and its proteinaceous inhibitor: a review. Carbohydr Res. 2010;345: 2583–2595. doi:10.1016/j.carres.2010.10.002

65. Anderson CT, Pelloux J. The Dynamics, Degradation, and Afterlives of Pectins: Influences on Cell Wall Assembly and Structure, Plant Development and Physiology, Agronomy, and Biotechnology. Annu Rev Plant Biol. 2025;76: 85–113. doi:10.1146/annurev-arplant-083023-034055

66. Chandel V, Biswas D, Roy S, Vaidya D, Verma A, Gupta A. Current Advancements in Pectin: Extraction, Properties and Multifunctional Applications. Foods. 2022;11: 2683. doi:10.3390/foods11172683

67. Wu H-C, Bulgakov VP, Jinn T-L. Pectin Methylesterases: Cell Wall Remodeling Proteins Are Required for Plant Response to Heat Stress. Front Plant Sci. 2018;9. doi:10.3389/fpls.2018.01612

68. Hyodo H, Terao A, Furukawa J, Sakamoto N, Yurimoto H, Satoh S, et al. Tissue Specific Localization of Pectin–Ca2+ Cross-Linkages and Pectin Methyl-Esterification during Fruit Ripening in Tomato (Solanum lycopersicum). PLoS One. 2013;8: e78949. doi:10.1371/journal.pone.0078949

69. Lara-Espinoza C, Carvajal-Millán E, Balandrán-Quintana R, López-Franco Y, Rascón-Chu A. Pectin and Pectin-Based Composite Materials: Beyond Food Texture. Molecules. 2018;23: 942. doi:10.3390/molecules23040942

70. Jia K, Wang W, Zhang Q, Jia W. Cell Wall Integrity Signaling in Fruit Ripening. Int J Mol Sci. 2023;24: 4054. doi:10.3390/ijms24044054

71. Steinwand BJ, Kieber JJ. The Role of Receptor-Like Kinases in Regulating Cell Wall Function. Plant Physiol. 2010;153: 479–484. doi:10.1104/pp.110.155887

72. Liu C, Yu H, Voxeur A, Rao X, Dixon RA. FERONIA and wall-associated kinases coordinate defense induced by lignin modification in plant cell walls. Sci Adv. 2023;9. doi:10.1126/sciadv.adf7714

73. Yao X, Humphries J, Johnson KL, Chen J, Ma Y. Function of WAKs in Regulating Cell Wall Development and Responses to Abiotic Stress. Plants. 2025;14: 343. doi:10.3390/plants14030343

